# Phytoplankton Photophysiology Reveals Depth Specific Zooplankton Grazing

**DOI:** 10.1101/2024.11.19.624385

**Authors:** Jason R. Graff, Amy E. Maas

**Author notes:** corresponding author, Jason R. Graff. **Author Contribution Statement:** JG and AM conceived of and performed the sample collection and experiments at sea. JG performed the data analysis and JG and AM co-wrote the manuscript. **Data Availability:** Data for this study can be found in NASA’s SeaWiFS Bio-optical Archive and Storage System (SeaBASS) (Figures 1 and 2) under the NASA EXPORTS Project and at https://github.com/JasonRGraff/The-Phytoplankton-Ecophysiology-Collaboration (Figures 3 and 4).

## Abstract

Marine particle forensics frequently uncover information on composition, age, size, and ecological history. Zooplankton fecal pellets are also studied for process-related data, such as grazing rates and carbon sequestration potential. Here, flow cytometric analyses of fecal pellet contents revealed intact phytoplankton with photophysiological characteristics mirroring those of free-living cells. Mapping the cytometrically derived properties of cells inside fecal pellets onto vertical profiles from free-living cells revealed the potential to estimate depth specific grazing by individual zooplankton. An experiment conducted at sea confirmed that the photophysiological characteristics of free-living phytoplankton from multiple depths, consumed by zooplankton, and excreted within fecal pellets are retained for at least 24 hours after grazing is initiated. These results have implications for high resolution modeling of individual or group specific zooplankton grazing dynamics that are critical for accurately linking zooplankton grazing in the surface ocean with the mesopelagic and deep ocean food webs and carbon export.

**Scientific Significance Statement:** Fecal pellet forensics have provided significant contributions to the study of zooplankton grazing and the marine carbon cycle. Gaps in knowledge about these processes remain, and continued investigations into fecal pellet contents and their fate are important for assessing connections between the surface and deeper ocean ecosystems. We describe a study conducted in the North Atlantic in the Spring of 2021 using flow cytometry to investigate fecal pellets contents. Observations that intact phytoplankton within the pellets had similar photophysiological properties to the free-living community led to a series of sample collections and experiments which provided a path forward for determining depth specific grazing by zooplankton community members. Phytoplankton survival after passing through zooplankton guts and being packaged into fecal pellets, with their potential for release far below the surface mixed layer, support prior observations of healthy phytoplankton communities at depth and validate this mechanism for the rapid transport of freshly fixed carbon to deep ocean systems. The results should be of interest to plankton ecologists and carbon cycle scientists connecting surface and deep ocean ecosystems as application of this approach at a broader scale will provide opportunities for high resolution modeling of individual and group specific zooplankton behaviors.

## Introduction

Sinking marine particles, like zooplankton fecal pellets, are an important component of the biological carbon pump (Siegel et al. 2023). The packaging of microscopic surface ocean phytoplankton and other particles by grazers into larger denser pellets generates a flux of organic carbon into the deep ocean estimated to account for < 1 and up to 100% of carbon export (Turner 2015), depending on the local conditions, with the sinking flux of particles largely dominated globally by fecal material (∼85%) (Nowicki et al. 2022). While the sizes, shapes, and densities of these ballasted particles vary by grazer type and species, it is ultimately their payload which conveys their importance for biogeochemical cycles. To this end, fecal pellet analysis has focused on bulk properties like carbon content (Urban-Rich et al. 1998; Stamieszkin et al. 2021) or more specific aspects such as stable isotopes (Altabet and Small 1990; Doherty et al. 2021), plastics (Kvale et al. 2020), or on processes such as sinking rates (Small et al. 1979; Feinberg and Dam 1998) in order to understand and quantify the significance of these properties at larger scales. Although these properties provide increasingly detailed insights into the gravitational pump, there remains a great deal of uncertainty in how the zooplankton grazing food web is structured. Knowing how much grazing is occurring, at what depth, and on which phytoplankton types has been identified as a critical knowledge gap whose parameters may strongly shape pelagic ecosystems and their role in the biological pump (Rohr et al. 2024).

Phytoplankton specific properties within zooplankton guts and fecal pellets, often in the form of pigment fluorescence measurements, have been used for estimating zooplankton grazing rates for many decades (Downs and Lorenzen 1985; Schnetzer and Steinberg 2002). An interesting aspect of this approach is that it relies on active fluorescence within a fecal pellet, which originates from intact cells and active pigments that have not been macerated or digested to the point of complete degradation. In fact, viable phytoplankton have been found to be present in fecal pellets with the potential for resurrection and proliferation given the right set of conditions, including escape from encapsulation due to pellet breakage or resuspension within the euphotic zone (Jansen and Bathmann 2007). More so, the presence of phytoplankton well below the euphotic zone has been documented previously with observations of intact cells down to greater than 4000 m (Agustí et al. 2015; Guo et al. 2018).

Their presence in the mesopelagic realm (∼200 – 1000 m) can be attributed to processes such as gravitational sinking, physical mixing, and zooplankton vertical migration and defecation with subsequent liberation from fecal pellets at depth. One example of a mixing event observed in real time resulted in the displacement of surface ocean communities to > 250 m, leaving phytoplankton and other microbial community members and their associated carbon well below the mixed layer and euphotic zone following the systems rapid re-stratification (Graff and Behrenfeld 2018; Baetge et al. 2022). This exported material is subject to bacterial degradation and grazing by resident and migrating zooplankton, as well as continued sinking of this material to greater depths for long-term export, subsequently fueling deep ocean communities with a source of relatively recently fixed carbon.

Disentangling the sources and amount of fresh organic material found and consumed at depth would add highly specific, or at minimum a more nuanced understanding, of the under-constrained carbon cycle pathways connecting the surface and deep ocean. Additionally, an ability to assess zooplankton depth specific grazing at the genus or species level with fine vertical resolution would provide a path forward for accessing ecological and carbon cycle science in a novel yet highly specific way. During the 2021 EXport Processes in the Ocean from RemoTe Sensing North Atlantic (EXPORTS NA) field campaign near the Porcupine Abyssal Plain (PAP) monitoring station, sampling partnerships were born out of shared interests in the ocean’s great septic system and its contribution to carbon export. Fecal pellets collected by the zooplankton science team were commodity items distributed for collaborative analyses. One such analysis included the flow cytometric investigations of the phytoplankton found within mesozooplankton fecal pellets. Intact cells were identified and found to have cellular level parameters similar to vertically distinct free-living communities observed throughout the water column. Experiments confirmed that these parameters would be robust for at least a 24 h grazing period. The approach and results presented below offer a potential method for fecal pellet forensics that could identify where mesozooplankton are consuming phytoplankton within the water column, providing high resolution information on depth and organism specific zooplankton grazing patterns.

## Materials and Procedures

The 2021 EXPORTS NA field campaign consisted of multiple research vessels targeting a retentive eddy near the PAP study site. An overview of the 2021 field campaign and physical setting can be found in Johnson et al. (2024). Work specific to this study was conducted aboard the HMS James Cook and focused on the center of the eddy that had been identified prior to embarkation, while additional vessels and autonomous platforms surveyed a broader area around the eddy center. The focus here is on a relatively small subset of mesozooplankton fecal pellets collected from organisms after capture by vertical net tows or created through on-board experiments and flow cytometric analyses of the free-living and fecal pellet bound phytoplankton communities.

### Mesozooplankton Fecal Pellets

Mesozooplankton used in this study were collect using nighttime 100 m vertical net tows of a 200 μm mesh, 1 m ring net. Individuals captured in the net tows were placed into glass jars with 0.2 μm filtered seawater. Fecal pellets generated from these animals with pre-existing gut contents (i.e. material grazed *in situ*) were collected after 12 – 24 hours. The pellets were gently pipetted from the experimental chambers and transferred into pre-filtered (0.2 μm) artificial seawater (deionized water plus sodium chloride) in clean 5 mL polycarbonate tubes just prior to processing for flow cytometric analyses of contents (see below). Additional fecal pellets were generated though experiments with the pseudothecosomatous pteropod species *Cymbulia peronei*, which had been allowed to fully evacuate their gut contents (> 24 h) prior to the experiment. After gut clearance, individuals were placed into glass jars filled with whole seawater collected from specific depths. After 24 hours of grazing in the dark in an incubation chamber set to approximate the *in situ* seawater temperature, fecal pellets that were generated from this known community of phytoplankton were similarly isolated into filtered artificial seawater prior to processing and analysis.

### Flow Cytometric Analysis of Free-living and Fecal Pellet Bound Phytoplankton

Whole seawater was collected multiple times per day from discrete depths using 10 L Niskin bottles on a CTD rosette. Samples for flow cytometric analyses were collected directly into clean 5 mL polycarbonate tubes and analyzed within 30 minutes of collection on a Becton Dickinson Influx Cell Sorter (ICS) or a Guava easyCyte 5HT flow cytometer (Guava). The ICS has a single blue (488 nm) laser with a small particle detector for forward scatter (FSC) detection with additional detectors for side scatter (SSC), and fluorescence at 530 and 692 nm (FL530 and FL692). The Guava was equipped with a 488 nm laser with detectors for FSC, SSC, red fluorescence at 695 nm (FL692) and yellow fluorescence at 583 nm (FL583). The signals from these detectors were used to discriminate *Synechococcus*, picoeukaryote, and nanoeukaryote groups. *Prochlorococcus* was not observed by flow cytometry and their absence was confirmed through high performance liquid chromatography analysis on samples collected throughout the field experiment (data not shown). The length of time for each sample analysis and sample flow rates determined at sea were used to calculate *in situ* concentrations of phytoplankton when using the ICS following Graff et al. (2018). Use of the Guava was limited to the shipboard feeding experiment and was not used to determine the *in situ* phytoplankton concentrations presented here.

Fecal pellet contents analyzed on the flow cytometers were liberated by vortexing the isolated pellets after they had been placed into the filtered artificial seawater. A subsample of the pre-filtered water containing the fecal pellet was collected as a control sample prior to vortexing. The control sample and a sample for the liberated contents were analyzed on the ICS or Guava within 10 minutes of sample creation.

Cytometry data was visualized and analyzed using FlowJo v10 to identify phytoplankton groups and counts and group-specific mean values of scattering and fluorescence parameters were exported for analysis. Here, the analysis was limited to the prokaryote *Synechococcus* as it was readily identified by the aforementioned parameters, was the most distinct group not obscured by other fluorescent particles in the fecal pellets, and is one of the most numerically abundant groups in the ocean.

## Results and Discussion

Free-living *Synechococcus* were observed from the surface ocean down to at least 500 m; concentrations were highest in the surface ocean with a significant attenuation in concentration beginning at approximately 50 m (Figure 1A). Figure 1B shows representative scattergrams of FSC versus FL692 from selected depth samples with arrows indicating the cluster of data associated with *Synechococcus*. Fewer cells are present in deeper samples (Figure 1A) and changes in the position of the clusters relative to the arrows (which had their placement on the plots held constant) are observed (Figure 1B). Shifts in the clusters or group mean values over depth or time have been linked to cell specific properties associated with the photophysiological state of phytoplankton groups (e.g. Graff et al. 2018). For example, when all other growth conditions are equal, cells experiencing higher growth irradiance have a lower FL per cell due to a reduction in pigments in high light environments.

**Figure 1.**
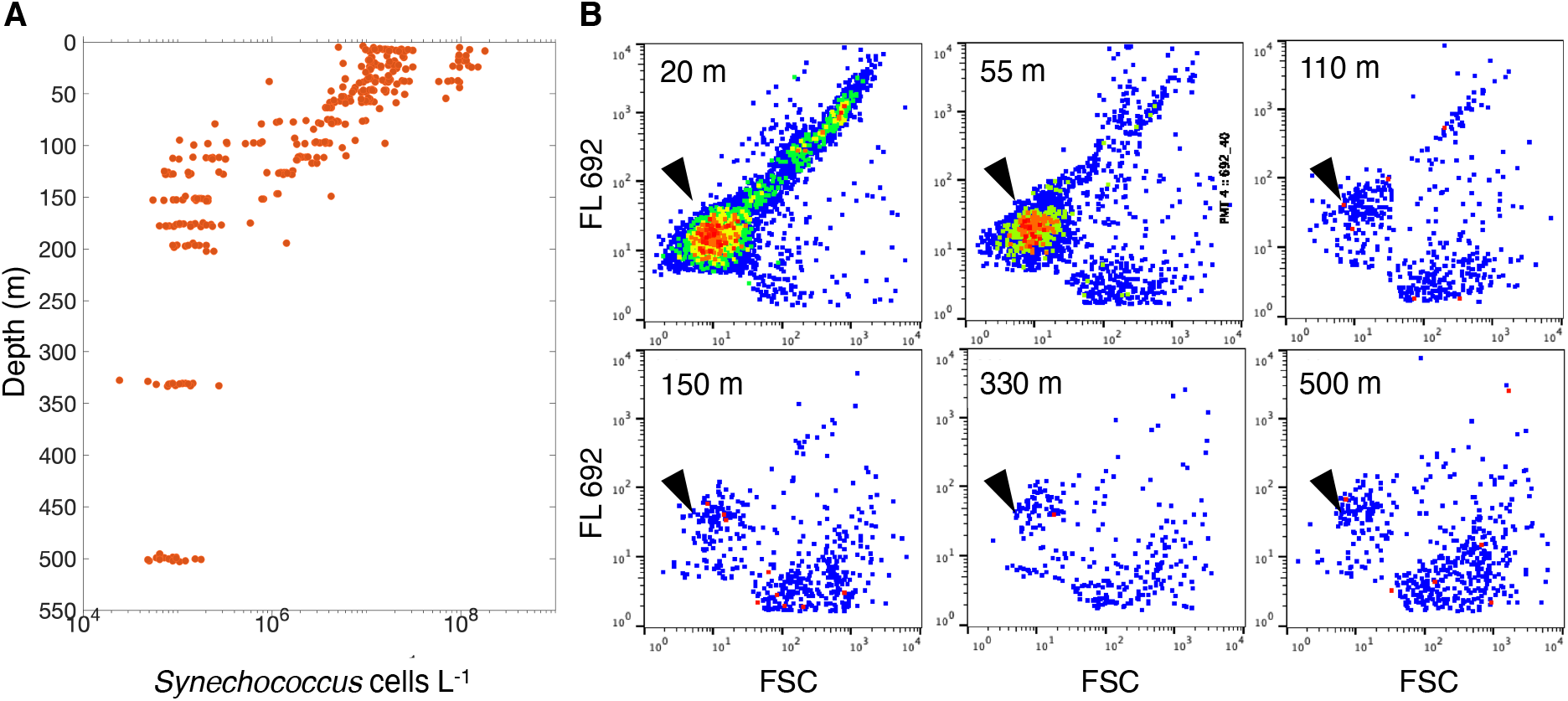
Flow cytometric analysis of phytoplankton communities during the NASA EXPORTS field campaign in the North Atlantic Ocean showing (**A)** the depth distribution of *Synechococcus* from 0 to 500 m for all Niskin collected samples analyzed on the ICS during the EXPORTS NA field expedition and **(B)** scattergrams of phytoplankton communities from selected depths of forward scattering (FSC) versus fluorescence at 692 nm (FL 692), arrows point to the cluster associated with *Synechococcus*.

Like berries in bear scat, phytoplankton compacted into fecal pellets were found within the supernatants of the disrupted pellets. Data from control samples are not shown here, but contained few if any phytoplankton. Figure 2 compares a sample of free-living phytoplankton collected at 35 m (Figure 2A) with liberated cells from disrupted fecal pellets of three representatives of the mesozooplankton collected (Figure 2B); salp, *Clio recurva*, and *Clio pyramidata*. Of note, multiple groups can be discerned within the fecal pellet material, but there is a high amount of fluorescent detrital material, which can be seen rising up from the origin of the Figure 2B panels, which complicates the ability to isolate and analyze all of the phytoplankton groups. Hence, the focus here is on *Synechococcus*, which due to its phycoerythrin content, gives it a higher fluorescence signal that can be detected on the ICS and Guava at 530 at 583 nm, respectively. While this method is not yet quantitative, the ratio of different cell types within the fecal pellets appears to be different either due to where cells in this pellet were consumed in the water column, selective ingestion by the zooplankton taxa under investigation, or resistance to gut processes. Nevertheless, it is clear that intact phytoplankton within sinking zooplankton fecal pellets can offer a rich source of recently fixed carbon to the deep ocean.

**Figure 2.**
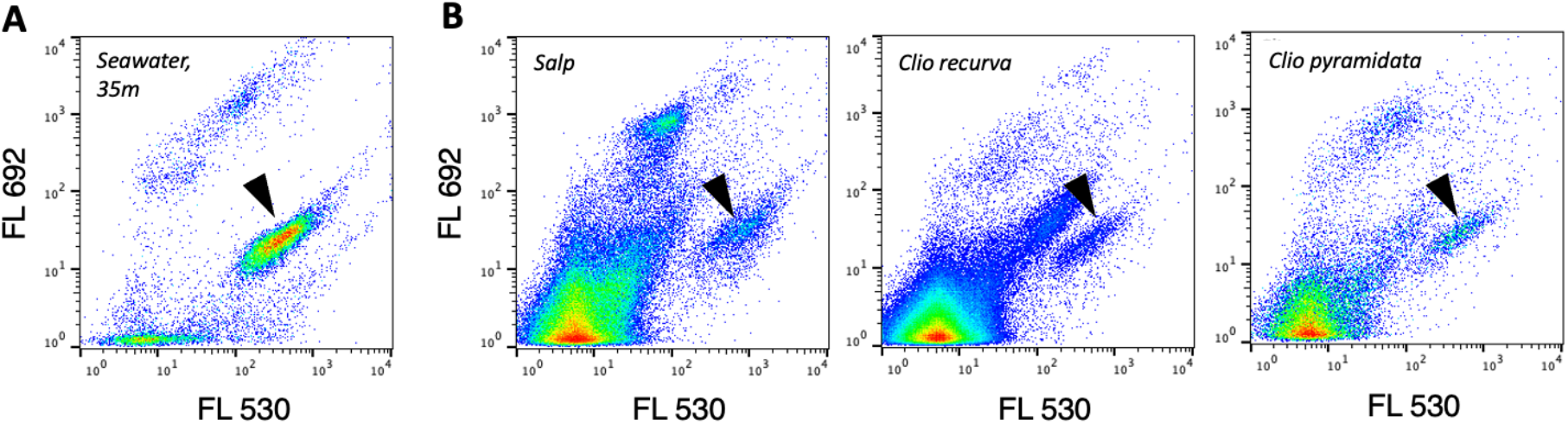
Flow cytometric analysis showing the scattergrams of fluorescence at 530 and 692 nm (FL530 versus FL 692) for **(A)** a representative phytoplankton community collected at 35 m during the NASA EXPORTS field campaign in the North Atlantic Ocean and **(B)** liberated particles and cells from fecal pellets from three representatives of the mesozooplankton community collected in 100 m vertical net tows. Arrows point to the cluster associated with *Synechococcus*.

So, are fecal pellets also harbingers of the free-”living” phytoplankton found at great depths? It seems that it is indeed one of multiple mechanisms. It is easy to envision a rich matrix of undigested living cells and recycled material within a fecal pellet, that if disrupted in the euphotic zone, would provide a near field pulse of nutrients for phytoplankton that had been trapped within it. A “grazer’s garden” of sorts. Disruption or degradation of the pellets below the euphotic zone and the phytoplankton’s fate is surely sealed in the absence of a deep mixing event to bring it back to the sunlit surface waters. Thus, a rapidly sinking and unmolested fecal pellet could shuttle “fresh” phytoplankton to great depths. The provenance of phytoplankton communities observed at depth is yet to be fully unraveled; however, there is more to this story which offers a potential path forward for high-resolution depth-resolved grazing.

The observation that the fluorescence and scattering parameters of phytoplankton found within fecal pellets are similar to free-living cells (Figure 2, compare *Synechococcus* in A and B panels) offers an ability to map these cells onto the depth resolved photophysiological state of the free-living *Synechococcus* using the ratio of FL530 to FSC (Graff and Behrenfeld 2018) (Figure 3). Figure 3A shows the depth profile of FL530:FSC for free-living *Synechococcus* from three Niskin rosette casts conducted on a single day. The model fit to the data (Figure 3A, dashed line and equation) was then applied to the FL530:FSC of the liberated cells from fecal pellets collected from 5 animals on the same day to determine the depth at which these cells may have been consumed (Figure 3B, blue diamonds). The range of depths where phytoplankton were grazed spans nearly 60 m and all samples mapped within the vertical range of the 100 m net tow. The net tow, however, cannot discern at what depth animals were when collected or where they were grazing. Is it possible that more specific information can be gleaned about the specific fine scale distribution and grazing habits of the zooplankton community directly from the fecal contents?

**Figure 3.**
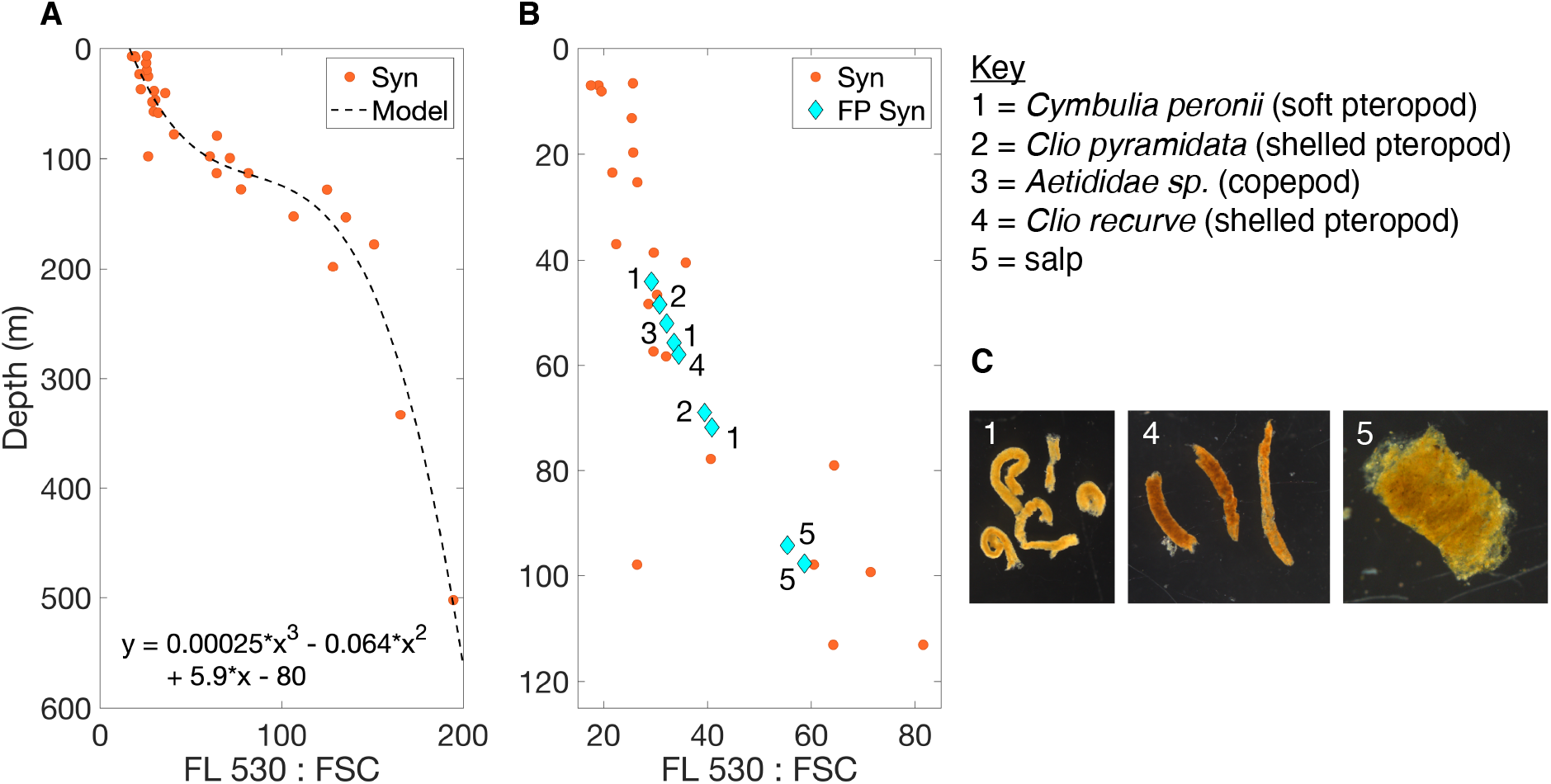
Depth resolved photophysiological state of phytoplankton based on flow cytometric analysis of fluorescence at 530 nm and forward scatter and a linear model describing that relationship for (**A**) *Synechococcus* from three CTD casts in the North Atlantic Ocean during the NASA EXPORTS 2021 field campaign and (**B**) the model applied to *Synechococcus* cell photophysiology recovered from 9 fecal pellets collected from animals within the same time frame to estimate grazing depth and relative to the free-living community. (**C**) Representative images of the densely packed fecal pellets collected, disrupted, and analyzed for phytoplankton community metrics.

While the cellular properties of phytoplankton in the fecal pellets resemble those of the free-living community, data from the natural gut content analysis does not resolve whether phytoplankton cells were altered through the process of ingestion and excretion and, thus, doubts remained on the depth-specific grazing implied in Figure 3B. Is the photophysiology of the ingested cells a bio-tracer preserved in the fecal pellets to the extent that the depth at which the cells were consumed can actually be determined? A simple ad hoc experiment was devised during the field expedition whereby a mesozooplankton, *Cymbulia peronii*, that had been starved for 24 hours in filtered seawater and had already evacuated its gut contents, were placed into whole seawater collected from specific depths. The photophysiological states of the phytoplankton (FL583:FSC) were recorded at the time the experiment began. After 24 hours of grazing, the newly formed fecal pellet contents were analyzed. Again, intact and apparently healthy phytoplankton were found in the fecal pellets. Aligning the photophysiological states of the liberated but recently ingested phytoplankton with those of the community collected from the *in situ* data offers an intriguing insight and path forward to realize depth-specific grazing by members of the zooplankton community (Figure 4). Free-living and pellet-derived *Synechococcus* had very similar FL583:FSC values (Figure 4, black versus pink data points for identical depths). Similar to Figure 3, a cubic polynomial was fit to the *in situ* profile data and an estimated depth value was calculated using the FL583:FSC ratio of the cells from the fecal pellets generated over the 24 h experiment. The minimum difference between the actual water collection depth and the depth estimated (Figure 4, blue diamonds) from the photophysiology of the fecal pellet cells was 1.5 m and the maximum difference was 10 m.

**Figure 4.**
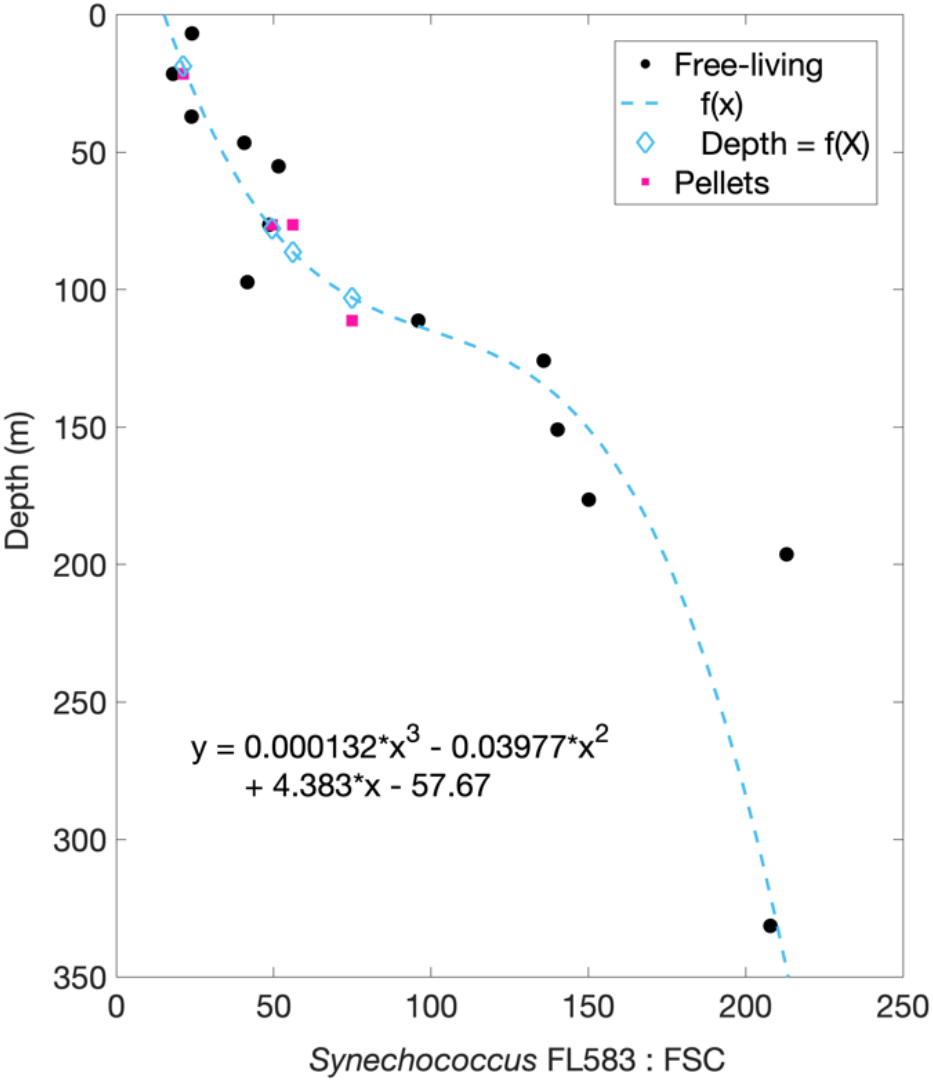
The ratio of flow cytometrically derived fluorescence at 583 nm (FL583) and forward scatter (FSC) for *Synechococcus* cells collected *in situ* (Free-living), cells liberated from fecal pellets (Pellets) generated during a 24 hour feeding experiment with the mesozooplankter *Cymbullia* sp. (pink; plotted at the depth where water was collected for the feeding experiment), and the depths estimated using a cubic polynomial fit to the *in situ* values (blue diamonds).

## Conclusions

The results of this concise ad hoc study offer a path forward for leveraging phytoplankton photophysiology to garner insights on zooplankton feeding habits and fecal pellet origination at very high vertical resolution. Further development and application of this approach could lead to establishing group or species-specific top-down impacts on phytoplankton communities in the surface ocean as well as new details for food web models linking zooplankton grazing with the deep ocean and carbon export. To achieve and realize such results, one can envision a series of fecal pellet collections from animals and sinking material and a series of grazing experiments focused on quantitative cytometric and biogeochemical measurements regarding fecal pellet contents. The fates and activities of apparently healthy phytoplankton within zooplankton fecal pellets remain elusive. Are they mostly doomed to enrich the deep ocean with their recently fixed carbon? Does proximity within the pellet foster high rates of gene transfer? Is there partisan survival across the encapsulated microbial groups and does liberation in the euphotic zone within in a nutrient rich pocket offer advantages to the survivors? These limited but fascinating observations provide intrigue and hope for approaches to continue to unravel the mysteries of the ocean’s great septic system.

## Acknowledgements

We would like to thank the captain and crew of the RRS James Cook and Discovery for their hard work and professionalism during the 2021 EXPORTS North Atlantic field campaign and to the EXPORTS leadership team for a successful expedition. This work was supported by funding from NASA Grants 80NSSC17KO568 supporting JG and 80NSSC17K0654 supporting AM.

